# Antimicrobial Resistance Surveillance Among Gram Negative Bacterial Isolates from Patients in Khartoum State Hospitals

**DOI:** 10.1101/486274

**Authors:** Hana Salaheldin Elbadawi, Kamal Mustafa Elhag, Elsheikh Mahgoub, Hisham N Altayb, Muzamil Mahdi Abdel Hamid

## Abstract

**Background:** Antimicrobial resistance (AMR) among Gram-negative bacilli is a global health problem. Surveillance of AMR is required to advise on empirical antimicrobial therapy. This study aimed at evaluating the frequency and the AMR patterns of Gram-negative isolates from patients treated in eight hospitals in Khartoum State, Sudan.

**Methods:** A cross-sectional laboratory based study was conducted over six months period at the microbiology department, Soba University Hospital, Khartoum State, Sudan. All Gram-negative isolates from blood, urine, wound, and sputum during the period of study were included.

**Results:** A total of 734 Gram-negative bacilli were isolated. *Klebsiella spp.* 249 (34%) was the most frequently encountered one, followed by *Pseudomonas spp.* 153(21%), *E.coli* 123(17%), *Acinetobacter spp.* 75 (10%), *Burkholderia cepacia* 42(6%), *Proteus spp.* 28(4%) *Enterobacter spp.* 28(4%), *Stenotrophomonas maltophilia* 21(2.8%), and others gram-negative bacilli 15(2.2%) The analysis of the antimicrobial susceptibility patterns showed that 134 (22.3%) isolates were multidrug resistant to three or more classes of antibiotics including cephalosporins, β-lactam-β-lactamase inhibitor group, quinolones, aminoglycosides and carbapenems.

**Conclusion:** This high level of resistance among Gram-negative bacilli in Khartoum state hospitals is alarming. The local health authorities are prompted to step up infection control program and introduce the concept of antimicrobial stewardship in Khartoum State hospitals.

## Introduction

Antimicrobial resistance (AMR) constitutes a continuously growing threat to the effective treatment of microbial infections [1]. The impact of AMR on human health, as well as the costs incurred on the health-care sector and the wider societal impact, are still largely unknown [2]. Antibacterial drugs are widely used worldwide both in human health and food industry. Overuse of these medications can favor the selection and the spread of multidrug resistant (MDR) bacteria [1]. Multi drug resistance is defined as resistance to at least three different antibiotic groups, as reported Masgala [3]. Antibiotic resistance among a variety of bacterial species is increasing in both healthcare settings as well as community one. Extended-spectrum β-lactamase and carbapenemase production are the most frequently emerging resistance mechanisms among Gram-negative bacilli [4]. Gram-negative bacilli including *Enterobacteriaceae* and non-lactose fermenting bacteria such as *Pseudomonas spp.* and *Acinetobacter spp.* are the main causes of hospital-acquired infection in critical care units [2,5]. The rate of antibiotic resistance among these pathogens has accelerated dramatically in recent years and has reached a pandemic scale [2]. According to the Centre for Disease Control and Prevention Gram-negative bacilli possess multiple modes of antibiotic resistance and are highly efficient in horizontally transferring resistance genes between species [6]. This problem of antimicrobial resistance is particularly pressing in developing countries, where the infectious disease burden is high and cost constraints prevent the widespread application of newer, more expensive agents [7].

AMR surveillance is the most important tool for assessing the burden of AMR and for providing the necessary antibiogram data, based on which the local, national, and global treatment strategies can be planned. Many surveillance studies on AMR are available in developed countries but unfortunately, studies on AMR surveillance are not adequate from developing ones. This surveillance study was undertaken in order to find out the different types of the AMR patterns of bacterial pathogens isolated from patients in Khartoum state, Sudan. This study may help in formulating antibiotic policies tailored to our hospitals. These data can be used as “information for action” antibiotic stewardship and interventions to optimize antibiotic prescribing practice, therefore prolongs the usefulness of existing antibiotics.

## Material and Methods

### Study design and clinical strains

This is a cross-sectional laboratory based study carried out in the department of medical microbiology Soba University Hospital (SUH), Sudan. A total of 734 Gram-negative isolates from patients treated in eight hospitals in Khartoum state, Sudan between October 2016 to February 2017. The isolates were collected from different clinical specimens including blood 243(33.1%), urine 230(31.3%), wounds 183(25%), sputum 22(3%), catheter tips 25(3.4%) and different body fluids 31(4.2%) (including CSF, peritoneal fluid, pleural fluid, acetic fluid, and synovial fluid). Microorganisms were grown on to Blood, Chocolate and MacConkey agar. Then, they were identified according to standard microbiological procedures (based on colony morphology, microscopy, and biochemical tests) [8]. Quality control strains were used in biochemical tests and antimicrobial susceptibility testing [*Escherichia coli* (ATCC #25922) and *Pseudomonas aeruginosa* (ATCC #27853)].

### Antimicrobial Susceptibility Testing

Susceptibility testing was performed by Kirby-Bauer disc-diffusion method for all isolates against the following antibiotic disc (Mast Diagnostic): Amoxycillin clavulanate (AMC) (30μg); Cefuroxime (CXM)(30μg); Cephalexin (CL)(30μg); Ceftriaxone (CRO)(30μg); Ceftazidime (CAZ) (30μg); Meropenem (MEM)(10μg); Imipenem (IPM) (10μg); Amikacin (AK) (30μg); Gentamicin (Gen)(10 μg); Ciprofloxacin (CIP)(5 μg); Trimethoprim-sulfamethoxazole (SXT) (25 μg); Temocillin (TEM) (30 μg); Azteroname (AZT)(30 μg); Nitrofrantoine (NIT) (300 μg). Results were interpreted according to the Clinical Laboratory Standards Institute (CLSI) guidelines [8].

### Classification of MDR Gram-Negative Bacilli

MDR has been considered for clinically significant Gram-negative bacilli (GNB) such as *Escherichia coli, Klebsiella spp., Pseudomonas spp.* and *Acinetobacter spp*., based on the above mentioned antimicrobial resistance definition. Classes of antibiotics used for MDR-GNB analysis were Aminoglycoside (AMG), Cephalosporins (CEPH), Carbapenems (CARB), and Fluroquinolones (FQ) as follows: bacteria that were MDR for 4 classes of antibiotics (AMG+ CEPH+ CARB+ FQ) and bacteria that were MDR for 3 classes of antibiotics (either AMG+CEPH +FQ, CARB+CEPH+FQ, AMG+CEPH+CARB, or AMG+FQ+CARB) [3].

Cephalosporins resistance was defined as resistance to ceftriaxone and ceftazidime except for *Pseudomonas species*, where only ceftazidime was used. Carbapenem resistance was defined as resistance to both meropenem and imipenem. Aminoglycoside resistance was defined as resistance to both gentamicin and amikacin, Ciprofloxacin resistance was considered an indication to fluoroquinolones resistance.

### Statistical analysis

Data were analysed using Microsoft Excel and SPSS version 20.0. Cross tabulation was used to present the different relations between data, qualitative data were performed through χ2test, and significance was set at *p*≤ 0.05. which performed to find the differences between bacterial isolates with resistance to at least one class of antibiotics by specimens (blood, urine, wound and other samples) *P*-values were determined for primary and secondary outcomes.

### Ethical consideration

Formal permission was obtained from the managers of Soba University Hospital and the Institutional Research Ethics Committee of the Institute of Endemic Diseases, University of Khartoum, approved this study under reference number 12/2017.

## Results

### Bacterial identification

Isolated Gram-negative bacilli showed different strains, including *E.coli* 123 (17%), *Klebsiella spp*. 249 (34%), *Pseudomonas spp*. 153 *(21%), Acinetobacter spp*. 75 (10%), *Burkholderia cepacia* 42 (6%), *Proteus spp*. 28 (4%) *Enterobacter spp*. 28 (4%), *Stenotrophomonas maltophilia* 21 (2.8%) and others gram-negative bacilli 15 (2.2%).

While isolates were distributed among the different hospital units, most of the pathogenic strains were isolated from neonatal intensive care unit (NICU) (42.7%) and pediatric units in (23.7%). *Klebsiella spp*. was the most isolated organism from all hospital units (Table 1).

**Table 1:** Distribution of Gram-negative bacilli among different hospital wards between October 2016 to February 2017

| Hospital ward | ICU | NICU | Medicine | Surgery | Renal Unit | Paediatric | P-value |
| --- | --- | --- | --- | --- | --- | --- | --- |
| Bacteria | N (%) | N (%) | N (%) | N (%) | N (%) | N (%) |  |
| *Klebsiella* spp. ((249) | 19 (39) | 77(42.3) | 45(30.6) | 34(33) | 24(31) | 50(29) | <0.001 |
| *E.coli*(123) | 2(12.2) | 9(5) | 45(30.6) | 22(21.3) | 16(21) | 25(14.2) | <0.001 |
| *Pseudomonas* spp. (153) | 9(18.4) | 40(21.9) | 24(16.3) | 17(16.5) | 19(24.3) | 44(25.1) | <0.001 |
| *Acintobacter* spp. (75) | 9(18.4) | 18(9.9) | 9(6.1) | 8(8) | 7(9) | 24(14) | <0.001 |
| *Burkholderia cepacia* (42) | 3(6.1) | 11(6) | 7(4.7) | 6(6) | 4(5.1) | 11(6.3) | <0.001 |
| *Proteus* spp. (28) | 1(2) | 3(2) | 4(2.7) | 9(9) | 3(4) | 8(5) | <0.001 |
| *Enterobacer* spp. (28) | 1(2) | 10(5.5) | 4(3) | 3(3) | 4(5.1) | 6(3.4) | <0.001 |
| *Stenotrophomonas* spp. (21) | 1(2) | 10(5.5) | 3(2) | 2(2) | 1(1) | 4(2) | <0.001 |
| Other Gram negative bacilli (15) | 0(0) | 4(2.2) | 6(4) | 2(2) | 0(0) | 3(2) | <0.001 |
| Total (734) | 49(7) | 182(42.7) | 147(20) | 103(14) | 78(11) | 175(23.3) | <0.001 |

(Other Gram-negative bacilli include *Citrobacter* species, *Serratia* species, *Vebrio vurneficus* and *Morganella morganii*)

With regard to the distribution of the isolates among different clinical specimens, *Klebsiella spp*. and *Pseudomonas spp*. were isolated mainly in blood specimens 39% and 25% respectively, while *Klebsiella spp*. and *E.coli* were 36% and 30% of urine samples. *Acinetobacter spp*. was mostly isolated from 25% of wound specimens (Table 2).

**Table 2:** Distribution of Gram-negative bacilli among different clinical samples between October 2016 to February 2017

| Specimen | Blood | Urine | Wounds | Sputum | Catheter<br>Tips | Body<br>Fluids | P-value |
| --- | --- | --- | --- | --- | --- | --- | --- |
| Bacteria | N (%) | N (%) | N (%) | N (%) | N (%) | N (%) |  |
| *Klebsiella spp.*<br>(249) | 94(39) | 82(36) | 50(27) | 7(32) | 7(28) | 9(29) | <0.001 |
| *E.coli*(123) | 15(6.1) | 70(30) | 31(17) | 2(9) | 1(4) | 4(12.9) | <0.001 |
| *Pseudomonas spp.*<br>(153) | 60(25) | 32(14) | 42(23) | 6(27.2) | 7(28) | 7(25) | <0.001 |
| *Acinetobacter spp.*<br>(75) | 18(7.4) | 13(5.6) | 25(14) | 6(27.2) | 7(28) | 2(25) | <0.001 |
| *Burkholderia*<br>*cepacia* (42) | 20(8) | 9(4) | 9(5) | 1(5) | 1(4) | 2(6.4) | <0.001 |
| *Proteus spp.* (28) | 6(2.5) | 7(3) | 12(6.5) | 0(0) | 2(8) | 1(3.2) | <0.001 |
| *Enterobacter spp.*<br>(28) | 13(5) | 8(3.5) | 7(4) | 0(0) | 0(0) | 0(0) | <0.001 |
| *Stenotrophomonas*<br>*spp.* (21) | 12(5) | 2(0.9) | 5(3) | 0(0) | 0(0) | 2(6.4) | <0.001 |
| Other Gram<br>negative bacilli<br>(15) | 5(2) | 7(3) | 2(1) | 0(0) | 0(0) | 0(0) | <0.001 |
| Total (734) | 243(33.1) | 230(31.3) | 183(25) | 22(3) | 25(3.4) | 32(4.2) | <0.001 |

(Other Gram-negative bacilli include *Citrobacter species*, *Serratia species*, *Vibrio vulnificus* and *Morganella morganii*)
Body fluids include (CSF, peritoneal fluid, pleural fluid, ascitic fluid, and synovial fluid)

### Antimicrobial Resistance Pattern of Clinical Isolates

Antibiotic resistance pattern are shown in Fig 1. Out of 734 isolates tested by disk diffusion test, the highest percentage of resistance 97%, and 93.5% were found against ampicillin and cephalexin, respectively, followed by amoxicillin/clavulanic acid 90%, cefotaxime 89.7%, ceftriaxone 88.4%, and ceftazidime 79.2%. In addition, co-trimoxazole and nitrofurantoin resistance were detected in 74.4 and 75.2 of isolates, respectively. Resistance rates also were high in ciprofloxacin 45.2%, gentamicin 52.5% and amikacin 18.3%. Meropenem and imipenem were the most effective antibiotic tested, resistant were observed with 21.6% and 16.2% of isolates, respectively.

**Fig 1:**
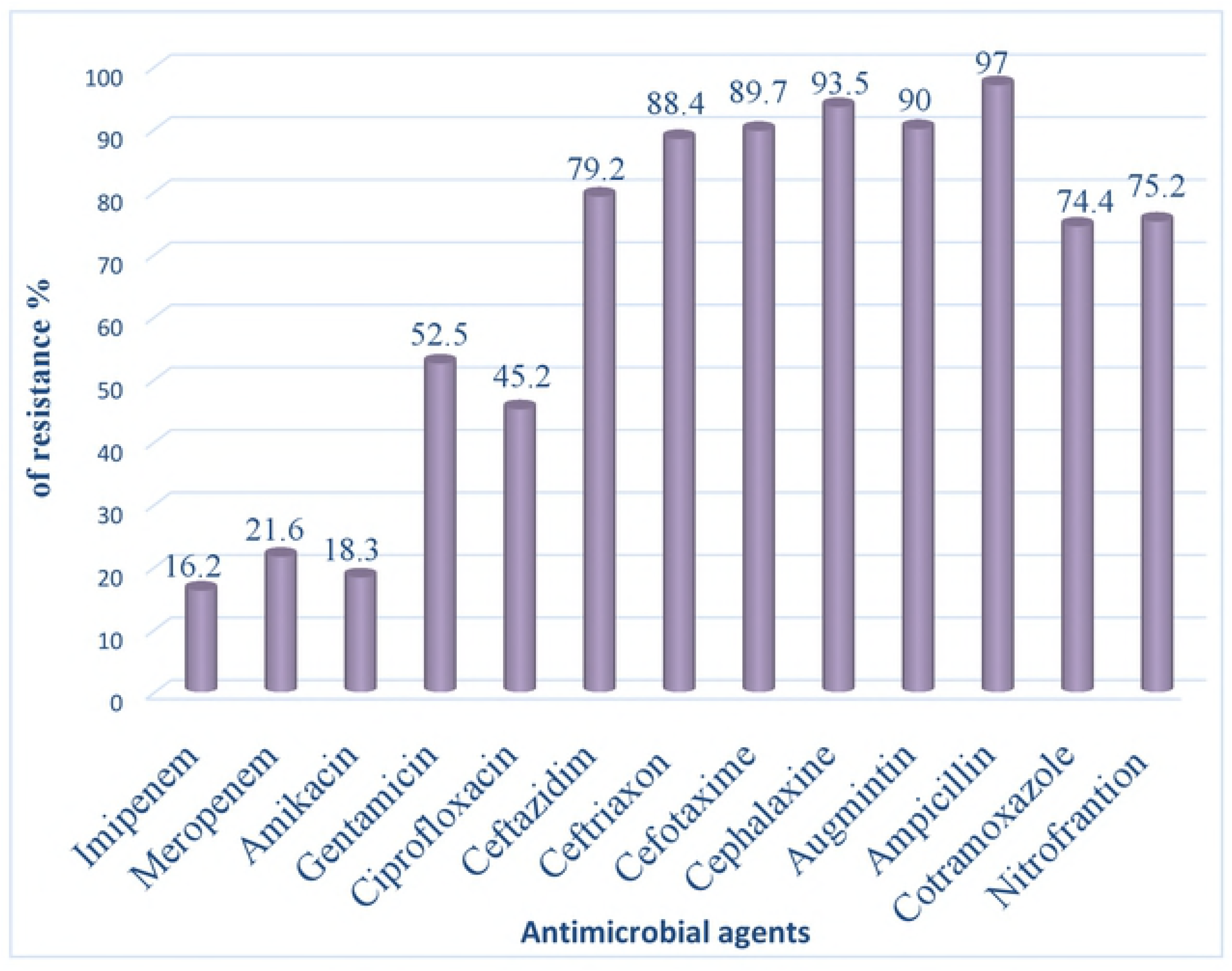
Antimicrobial Resistance pattern among different Gram-negative bacilli isolated from patients treated at Khartoum state hospitals between October 2016 to February 2017.

The antimicrobial resistance patterns of most commonly isolated organisms are shown in Fig 2. The analysis of the antimicrobial susceptibility patterns of the study isolates showed high rate of MDR organisms that were resistant to three or more classes of antibiotics, including carbapenem and aminoglycosides. This pattern mainly among *Acinetobacter spp., Pseudomonas spp*., and *Klebsiella spp*.

**Fig 2:**
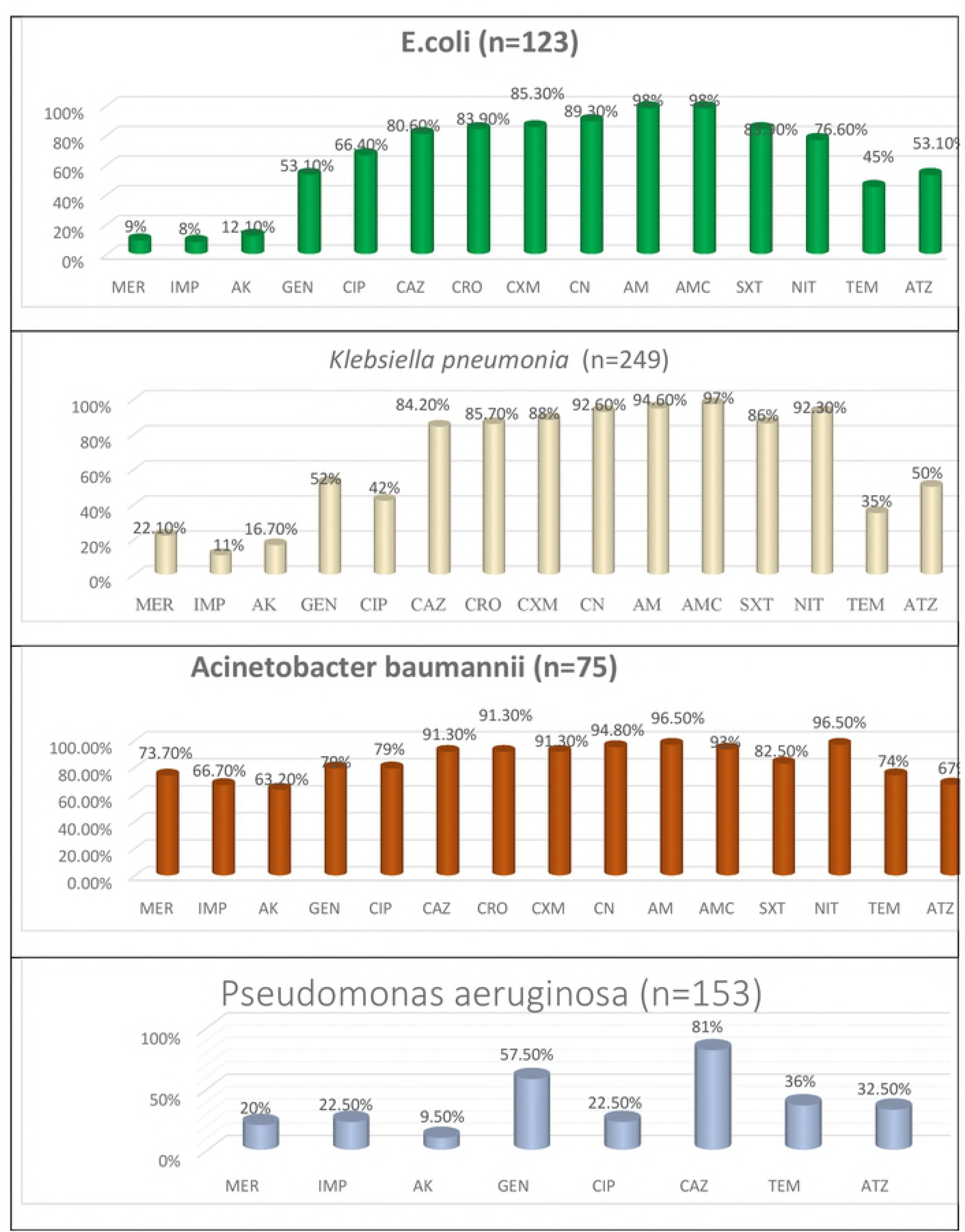
Sensitivity pattern among commonly isolated organisms’ different antibiotics between October 2016 to February 2017. (MER-Mcropcnem, IMP-Imipcncm.AK-Amikacin. GEN-Gcntamicin.CIP-Cipi?f?oxacin.CAZ-Ccftazidimc, CRO-Ccftria\oncCXM-Ccfuroximc. CN-ccphalexin.

The most clinical isolates in Enterobacteriaceae family were *E.coli* and *Klebsiella spp*. in both of these GNB, high rate of resistance was observed against quinolones, cephalosporins and β-lactam group of drug. Resistance to carbapenem was also statistically significantly high.

### Multidrug Resistance in Gram Negative Bacilli (Co Resistance Patterns)

The multidrug resistance pattern among clinical isolates of gram-negative bacilli have been shown in Table 3. Of 600 GNB isolates recovered, 134 (22.3%) isolates were MDR i.e., resistant to at least three or more classes of antimicrobial agents. About 48(8%) of all Gram negative isolates were 4MDR (resistant to four classes of antimicrobial drug), while 86 (14.2%) were 3MDR (resistant to three classes of antimicrobial drug). It was observed that 50% of *Acinetobacter spp*. were resistant to four major classes of antibiotics. Significant resistance to carbapenem was noted among all gram-negative bacteria.

**Table 3:** Multi drug resistance pattern among gram-negative isolates between October 2016 to February 2017

| Class | Resistance to | *Klebsiella*<br>*spp.* (249) | *E.coli*<br>(123) | *Pseudomonas*<br>*spp.* (153) | *Acinetobacter*<br>*spp.* (75) | Total<br>(600) |
| --- | --- | --- | --- | --- | --- | --- |
| 4MDR | AMG+ CEPH+<br>FQ+ CARB | 9 (3.6%) | 1 (0.8%) | 0 (0%) | 38(50.6%) | 48 (8%) |
| 3MDR | AMG+ CEPH+<br>FQ | 31(12.4%) | 5(4%) | 4 (2.6%) | 2 (2.6%) | 42 (7%) |
| 3MDR | CARB+<br>CEPH+ FQ | 6 (2.4%) | 0 (0%) | 8 (5.2%) | 8 (10.6%) | 22(3.6%) |
| 3MDR | AMG+ CEPH+<br>CARB | 2 (0.8%) | 0 (0%) | 5 (3.2%) | 1 (1.3%) | 8 (1.3%) |
| 3MDR | AMG+ FQ+<br>CARB | 8 (3.2%) | 1(0.8%) | 4 (2.6%) | 1 (1.3%) | 14<br>(2.3%) |
| Total |  | 56(22.4%) | 7 (5.6%) | 21(13.7%) | 50 (66.6%) | 134<br>(22.3%) |

AMG = Aminoglycoside. CEPH= Cephalosporins, FQ= Florquinolones, CARB= Carbapenem.

## Discussion

Infection with MDR Gram-negative bacilli is a major problem worldwide, associated with increased patients morbidity and mortality [9]. In Sudan, the increasing number of MDR bacteria is a real clinical challenge [10,11]. This study was undertaken; to determine the occurrence of different types of AMR pattern of the bacterial pathogens isolated from patients treated in various wards of hospitals.

In this study, *Klebsiella* and *Pseudomonas* strains were more prevalent in blood specimens while *Klebsiella* and *E.coli* strains were more frequently isolated in urine specimens.

*Klebsiella pneumoniae* is an important causative agent of nosocomial and community-acquired Gram-negative bacteremia. It can cause various infections, including blood stream infections, wound infections, pneumonia, urinary tract infections and intra-abdominal infections [12,13]. In this study, *Klebsiella spp*. was the most pathogenic strain isolated from the blood specimens mainly in neonatal sepsis in a high rate (39.8%). Most of these strains resistant to cephalosporins and other class of antibiotics including carbapenem as reported worldwide [7,13,14].

*E.coli* is the commonest urinary tract pathogen causing complicated and uncomplicated UTI [16] but in this study most observed pathogens in urine specimens were *E.coli* 30% and *Klebsiella spp*. 36%. This finding in Sudan is concordant with that of de Francesco et al, 2007 where *E.coli* was found in 42.4% of Gram-negative isolates [17]. This also goes with results that were obtained in Tanzania where *E.coli* was detected in 38% of the Gram-negative isolates and 25% of all Isolates [18]. Likewise, many authors have the same finding in Pakistan and India [18,19].

Non-lactose Fermenting Gram negative bacilli such as *Pseudomonas spp*. and *Acinetobacter spp*. were the mostly isolated from ICU patients from blood, wound and sputum specimens, the isolation rates were 18.4% for both. In this study we found that *Pseudomonas spp*. were associated with 25% of blood stream infections and 23% of wound infection while *Acinetobacter spp*. mainly with wound infection in 25% concordant with Gales et al 2010 [20].

*Pseudomonas spp*. was reported as highly associated in health care setting by Jean-Louis Vincent et al.,[21]. While Javeri Jitendra R et al., also reported that *Acinetobacter spp*. as the second most common in an ICU of tertiary care center [22].

Resistance of gram negative bacilli has emerged widely and multidrug resistance has been reported by many studies causing challenge to treatment of nosocomial infections. The resistance pattern was commonly reported in classes such as cephalosporins, carbapenem, aminoglycosides and quinolones [2,23–25]. In this study, we observed high rate of ESBL, resistant to ceftazidime and ceftriaxone in addition to resistance to ampicillin and amoxicillin/clavulanic acid.

The most resistant strain was *Acinetobacter spp*., being resistant to all four classes of antibiotics used in 50.6% of isolates. About 73.7% of *Acinetobacter* were found to be resistant to meropenem, and 66.7% to imipenem, while in cephalosporins classes more than 91% of the isolates were resistant. *Acinetobacter spp*. also have high resistance rate to Aminoglycosides and quinolones 63.2% for amikacin and 79% for both gentamicin and ciprofloxacin. This increasing resistance among *Acinetobacter spp*. has become a public-health issue because they play an important roles in nosocomial infections [24].

*Klebsiella spp*. and *E.coli* were the most isolated pathogens in this study with relatively high resistance rate, about 9 to 22% of them were resistant to carbapenem. And they have highly resistant rate to ceftazidime and cephalexin, 80.6% and 92% respectively. This is much higher than the observation in 2013 by Ali in Soba University hospital that Ceftriaxone and ceftazidime resistance ranged from 56% to 79% [14,26]. Aminoglycoside resistance among *Klebsiella spp*. and *E.coli* was 16.7% and 12.1% respectively to amikacin, which is lower than Normratha 2015 observation who found the resistance rate to amikacin 37% of *Klebsiella spp*. and 23% of *E.coli* [23]. Gentamicin resistance among both *Klebsiella spp*. and *E.coli* was up to 53%, which is slightly higher than Normratha study [23]. *E.coli* was highly resistant to Quinolones group like ciprofloxacin in 66.4% than *Klebsiella spp*. 42% this finding is much lower than observation by Moolchandani 2017 [25].

*Pseudomonas spp*. was significantly resistant to carbapenem in 22% of isolates and was highly resistant to ceftazidime in 81% followed by gentamicin 57.5%, ciprofloxacin in 22.5% and amikacin 9.5%. That was reported in many studies [4,27].

The high level of resistance in the current study can be attributed to the unrestricted use of antibiotics in Sudanese hospitals, which plays an important role in increasing carbapenem resistance [25]. During this study, 134 gram negative bacilli resistance to three or four classes of antibiotics in a period of six months were isolated, which is relatively higher rate than study in the period performed in SUH from January 2011 to June 2013 and reported 80 bacterial strains resistant to all available antibiotics including meropenem [26].

In Sudan most laboratories do not test for ESBL and Carbapenemase producer and report; based on disc diffusion test; an ESBL producer as sensitive to cephalosporins. ESBL producer in addition to cephalosporin resistant it may be associated with other resistance gene like *qnr* of quinolones [28]. This may in turn give false impression to the clinicians and mask the true picture of the high prevalence of antibiotic resistance. Moreover, there is limited choice of available antibiotics other than cephalosporins in Sudan that increased the prescription of cephalosporin for treatment of infectious diseases and meropenem for ESBL producer.

## Conclusion

In conclusion, there was high prevalence of Gram-negative bacterial pathogen associated hospital and community acquired infections with increasing resistance to available antibiotics. We need to implement of strict infection control measures and activate the antimicrobial stewardship, policed to decrease the spread of MDR pathogens in Sudanese hospitals.

## Acknowledgments

We would like to thank the technical staff of Medical Microbiology Department in Soba University Hospital, University of Khartoum for their help in strain and data collection. This research received partial research fund from the Ministry of Higher Education and Scientific Research, Sudan.

## Transparency Declaration

No conflict of interests to declare.

## References

1. Harris P, Paterson, D and Rogers B. Facing the challenge of multidrug-resistant gram-negative bacilli in Australia. Clin Focus. 2015;202:243–7.

2. Mehrad B, Clark NM, Zhanel GG, Iii JPL. Antimicrobial Resistance in Hospital-Acquired Gram-Negative Bacterial Infections. Chest. 2015;1413–21.

3. Masgala A, Kostaki K II. Multi Drug Resistant Gram Negative Pathogens in Long Term Care Facilities: A Steadily Arising Problem. J Infect Dis Diagn. 2015;(1):101.

4. Karaiskos I, Giamarellou H. Multidrug-resistant and extensively drug-resistant Gram-negative pathogens: current and emerging therapeutic approaches. Expert Opin Pharmacother. 2014;15(10):1351–70.

5. Rosenthal VD, George D, Mehta Y, Leblebicioglu H, Ahmed Z, Al-mousa HH, et al. International Nosocomial Infection Control Consortiu (INICC) report, data summary of 43 countries for 2007–2012. Device-associated module. Am J Infect Control. 2014;42:942–56.

6. Huang TD, Bogaerts P, Berhin C, Hoebeke M, Bauraing C, Glupczynski Y. Increasing proportion of carbapenemase-producing Enterobacteriaceae and emergence of a MCR-1 producer through a multicentric study among hospital-based and private laboratories in Belgium from September to November 2015. Eurosurveillance. 2017.

7. Report G. Antimicrobial resistance. 2014.

8. Testing S. M100 Performance Standards for Antimicrobial. 27th ed. 2017. 106–143 p.

9. Zilberberg MD, Nathanson BH, Sulham K, Fan W, Shorr AF. Multidrug resistance, inappropriate empiric therapy, and hospital mortality in Acinetobacter baumannii pneumonia and sepsis. Crit Care. 2016;20:1–10.

10. Yousif M. The prevalence of Extended Spectrum β-lactamase and AmpC-Producing Bacteria in a Sudanese Tertiary Hospital. Sudan Med J. 2015;5(3):10–17.

11. Ibrahim M, Bilal N, Hamid M. Comparison of phenotypic characteristics and antimicrobial resistance patterns of clinical Escherichia coli collected from two unrelated geographical areas. Glob J Heal Sci. 2014;(6):126–35.

12. Meatherall BL, Gregson D, Ross T, Pitout JD, & Laupland KB. Incidence, risk factors, and outcomes of Klebsiella pneumoniae bacteremia. Am J Med. 2009;122:866–873.

13. Cao X, Xu X, Zhang Z, Shen H, Chen J, Zhang K. Molecular characterization of clinical multidrug-resistant Klebsiella pneumoniae isolates. Ann Clin Microbiol Antimicrob. 2014;1:13–6.

14. Ali MA. the prevalence and characterization of antibiotic resistance among Gram-negative bacilli. University of Khartoum; 2013.

15. Tzouvelekis LS, Markogiannakis A, Psichogiou M, Tassios PT, & Daikos GL. Carbapenemases in Klebsiella pneumoniae and other Enterobacteriaceae: an evolving crisis of global dimensions. Clin Microbiol Rev. 2012;25:682–707.

16. Karlowsky J, Jones M, Thornsberry C, Critchley I, Kelly L, Sahm D. Prevalence of anti-microbial resistance among urinary tract pathogens isolated from female outpatients across the US in 1999. Int J Antimicrob Agents. 2001;18:121–7.

17. de Francesco MA, Giuseppe R, Laura P, Riccardo N NM. Urinary tract infections in Brescia, Italy: Etiology of uropathogens and antimicrobial resistance of common uropathogens. Med SciMonit. 2007;13:136–44.

18. Blomberg B, Olsen BE, Hinderaker SG, Langeland N, Gasheka P, Jureen R, Kvale G, and Midtvedt T. Antimicrobial resistance in urinary bacterial isolates from pregnant women in rural Tanzania: implications for republichealth. Scandinavian. J Infect Dis. 2005;37(4):262–8.

19. Haider G, Zehra N, Munir AA and HA. Risk factors of urinary tract infection in pregnancy. J Pak Med Assoc. 2010;60(3):21–36.

20. Zavascki AP, Carvalhaes CG, Picao RC GA. Multidrug-resistant Pseudomonas aeruginosa and Acinetobacter baumannii: resistance mechanisms and implications for therapy. Expart Rev Anti infet ther. 2010;(8):71–93.

21. Vincent JL, Rello J, Marshall J, Silva E, Anzueto A, Martin CD et al. International study of the prevalence and outcomes of infection in intensive care units. JAMA. 2009;302(21):2323–9.

22. Javeri JR, Patel SM, Nayak SN, Desai K PPA. study on bacteriological profile and drug sensitivity & resistance pattern of isolates of the patients admitted in intensive care units of a tertiary care hospital in Ahmadabad. Natl J Med Res. 2016;2(3):330–4.

23. Nandihal NW. Profile of Urinary Tract Infection and Quinolone Resistance among Escherichia coli and Klebsiella species isolates. IntJCurrMicrobiolAppSci. 2015;4(7):749–56.

24. Diene SM RJ. Carbapenemase genes and genetic platforms in Gram-negative bacilli: Enterobacteriaceae, Pseudomonas and Acinetobacter species. Clin Microbiol Infect Off Publ Eur Soc Clin Microbiol Infect Dis. 2014;30(9):831–8.

25. Moolchandani K, Sastry AS, Deepashree R, Sistla S, Harish BN, Mandal J. Antimicrobial Resistance Surveillance among Intensive Care Units of a Tertiary Care Hospital in Southern India. J Clin Diagn Res. 2017;11(2):1–7.

26. Elhag KM. Review Article Diversification of antibiotics as a means to control antimicrobial resistance and improve treatment options in Sudan. Sudan Med J. 2013;49(3):128–35.

27. Magiorakos A-P, Srinivasan A, Carey RB, Carmeli Y, Falagas ME, Giske CG, et al. Multidrug-resistant, extensively drug-resistant and pandrug-resistant bacteria: an international expert proposal for interim standard definitions for acquired resistance. Clin Microbiol Infect. 2012;18(3):268–81.

28. Paterson DL, Mulazimoglu L, Casellas JM, Ko W-C, Goossens H, Von Gottberg A, et al. Epidemiology of Ciprofloxacin Resistance and Its Relationship to Extended-Spectrum - Lactamase Production in Klebsiella pneumoniae Isolates Causing Bacteremia. Clin Infect Dis. 2000;30(3):473–8.

